# Comparative analysis of six adeno-associated viral vector serotypes in mouse inferior colliculus and cerebellum

**DOI:** 10.1101/2024.10.17.618966

**Authors:** Isabelle Witteveen, Timothy Balmer

**Affiliations:** School of Life Sciences, Arizona State University, Tempe, AZ, 85287, USA

## Abstract

Adeno-associated viral vector (AAV) serotypes vary in how effectively they express genes across different cell types and brain regions. Here we report a systematic comparison of the AAV serotypes 1, 2, 5, 8, 9, and the directed evolution derived AAVrg, in the inferior colliculus and cerebellum. The AAVs were identical apart from their different serotypes, each having a synapsin promotor and expressing GFP (AAV-hSyn-GFP). Identical titers and volumes were injected into the inferior colliculus and cerebellum of adult male and female mice and brains were sectioned and imaged 2 weeks later. Transduction efficacy, anterograde labeling of axonal projections, and retrograde labeling of somata, were characterized and compared across serotypes. Cell-type tropism was assessed by analyzing the morphology of the GFP-labeled neurons in the cerebellar cortex. In both the cerebellum and inferior colliculus, AAV1 expressed GFP in more cells, labeled a larger volume, and produced significantly brighter labeling than all other serotypes, indicating superior transgene expression. AAV1 labeled more Purkinje cells, unipolar brush cells, and molecular layer interneurons than the other serotypes, while AAV2 labeled a greater number of granule cells. These results provide guidelines for the use of AAVs as gene delivery tools in these regions.

**Significance:** AAVs have become ubiquitous gene expression tools in neuroscience research and are becoming more common in clinical settings. Naturally occurring and engineered serotypes have varying abilities to infect neurons and cause them to produce proteins of interest. The efficacy of AAV transduction in specific cell types depends on many factors and remains difficult to predict, so an empirical approach is often required to determine the best performing serotype in each population of cells. In the present study we show that AAV1 produces the highest expression in these two regions, labels the most axonal projections, and labels Purkinje cells and unipolar brush cells better than the other serotypes tested, while AAV2 labels granule cells most effectively.

## Introduction

Adeno-Associated Viruses (AAVs) have become critical tools in neuroscience research and hold substantial promise in clinical gene therapy applications. AAVs have been developed to treat some genetic disorders that result from impaired gene expression, including specific forms of inherited retinal disease and deafness (Haery et al., 2019; Wang et al., 2019; Haggerty et al., 2020; Xue et al., 2023; Lv et al., 2024). AAVs are small, ∼24 nm, viruses belonging to the dependoparvovirus genus, named for their inability to self-replicate without the presence of helper viruses such as Adeno virus (Atchison et al., 1965; Hoggan et al., 1966; Haery et al., 2019; Wang et al., 2019; Haggerty et al., 2020). Despite their prevalence in the human population–40-80% of humans are seropositive for antibodies against AAVs– they are not known to cause any human disease (Wang et al., 2019). Their low immunogenicity and ability to induce long-term gene expression in post-mitotic cells, have made AAVs attractive candidate vectors for use in gene therapy (Snyder and Moullier, 2011; Haery et al., 2019; Wang et al., 2019; Haggerty et al., 2020).

AAVs have become widely adopted tools for gene expression in neuroscience research. For example, AAVs are routinely used to express fluorescent reporters for the visualization and mapping of neuronal circuits. The delivery of Cre and other recombinases allows conditional gene expression or deletion to examine the functions of proteins, cell types, and circuits. The expression of optogenetic and chemogenetic tools can be used to control the activity of neurons. The delivery of calcium and voltage sensitive fluorescent indicators allows the visualization of neuronal activity. Many more tools are available and continue to be developed through the use of AAVs (Yizhar et al., 2011; Haery et al., 2019). Given their utility, analyses of the compatibility of different AAV serotypes in specific brain regions have become essential.

A variety of naturally occurring AAV serotypes have been found, each with varying capsid proteins and genomes (Snyder and Moullier, 2011; Wang et al., 2019). The effectiveness of AAV transduction is determined by interactions between capsid proteins and cell surface receptors, internalization, and subsequent downstream effects (Snyder and Moullier, 2011; Wang et al., 2019). Since serotypes recognize distinct receptors, this leads to species-specific, tissue-specific, and cell-type specific tropism (Snyder and Moullier, 2011; Wang et al., 2019). Pseudotyping takes advantage of this natural tropism by packaging the recombinant AAV genome into the capsids of different serotypes to increase transduction efficacy (Burger et al., 2004). Numerous studies have since characterized differences in AAV transduction between serotypes in various brain regions, including the nigrostriatal system, hippocampus, cerebral cortex, and a variety of other midbrain regions, utilizing both local and systemic injections (Wang et al., 2003; Burger et al., 2004; Paterna et al., 2004; Cearley and Wolfe, 2006; Taymans et al., 2007; Klein et al., 2008; Zincarelli et al., 2008; McFarland et al., 2009; Blits et al., 2010; Mason et al., 2010; Hutson et al., 2012; Aschauer et al., 2013; Holehonnur et al., 2014; Watakabe et al., 2015). These studies demonstrate the presence of region-specific tropism and underscore the necessity of empirical comparisons of AAV serotypes in well-controlled experiments. Quantitative analyses of the transduction efficacy of AAV serotypes in the inferior colliculus (IC) and cerebellum have not been reported.

The cerebellum is a structure necessary for coordinated movement and has been increasingly implicated in cognitive functions, while the IC is a major processing center of ascending and descending auditory pathways (Apps and Garwicz, 2005; Winer and Schreiner, 2005; Reeber et al., 2013; De Zeeuw et al., 2021). Previous research has utilized AAVs in these regions, but few have reported the effectiveness of different AAV serotypes. The transduction patterns of Purkinje cells have been examined for some serotypes (Kaemmerer et al., 2000; Hirai, 2008; Bosch et al., 2014; Kim et al., 2015; El-Shamayleh et al., 2017). Other studies have compared the transduction efficacy of some AAV serotypes at early postnatal ages (Broekman et al., 2006), differences in short-term synaptic plasticity following AAV-mediated channelrhodopsin expression (Jackman et al., 2014) or transduction patterns of an individual AAV serotype (Alisky et al., 2000). The goal of this study was to characterize the transduction and tropism of commercially available AAV serotypes following in vivo injection into the IC and cerebellum, which can inform further utilization of AAVs in these regions.

## Methods and Materials

### Animals

C57BL6/J mice of both sexes were used in this study (17 males, 15 females). The ages ranged from 2-3 months old. Mice were bred in a colony maintained in the animal facility managed by the Department of Animal Care and Technologies and all procedures were approved by [Author University] Institutional Animal Care and Use Committee under protocol #21-1817R.

### Adenoassociated Viruses

All viruses were purchased from Addgene containing EGFP under the control of the human synapsin promoter (Addgene plasmid # 50465; RRID: Addgene_50465). The serotypes included AAV1, AAV2, AAV5, AAV8, AAV9 and AAVrg. AAVrg was engineered to infect axons (Tervo et al., 2016), in addition to somata at the injection site. Images of labeling with AAVrg are shown in some figures, although this serotype was only included in the statistical comparisons of cell type tropism in the cerebellar cortex. Aliquots of 5 µL were stored at -80°C. Identical titers were used across all injections, 8*10^12^ GC/ml (4*10^9^ GC per 500 nl injection). Aliquots were thawed on ice and diluted with sterile saline to achieve this titer on the day of injection.

### Viral Injections

Viral injections were made into the cerebellum and the inferior colliculus under isoflurane anesthesia. Glass capillaries were pulled on a horizontal puller and beveled at a 45-degree angle to a 20-30 µm inside diameter. The glass pipettes were filled with mineral oil and placed on the injector (Nanoliter 2020, World Precision Instruments). Mice were weighed and then anesthetized in a knockdown chamber at a flow rate of 500 ml/min with 5% isoflurane, then placed in a stereotax with the flow rate reduced to 50 ml/min with 1-2% isoflurane and adjustments were made based on the frequency of respiration of the mouse using a digital vaporizer (SumnoSuite, Kent Scientific). Ophthalmic lubricant was applied to the eyes of the mouse to prevent drying. Following fur removal, the scalp was cleaned using 7.5% povidone iodine. An incision was made in the scalp and one or two small holes were drilled in the skull with a burr drill. Virus was placed on parafilm and then taken up into the pipette. The pipette was slowly lowered into the brain at a rate of approximately 10 µm/s. Three minutes after the pipette reached the injection site 500 nl of virus was injected at 5 nl/s. Three minutes after the injection the pipette was slowly retracted at a rate of 10 µm/s. Injections were made at the stereotaxic coordinates of 7.2 mm caudal, 0.5 mm lateral to bregma, and 2.9 mm ventral to the surface of the brain for the cerebellum and 5.5 mm caudal, 1.1 mm lateral to bregma, and 1.0 mm ventral to the surface of the brain for the right IC.

### Immunohistochemistry and Imaging

2 weeks after injection mice were overdosed with isoflurane and perfused through the heart with 0.01 M phosphate buffered saline, 7.4 pH (PBS) followed by 4% paraformaldehyde in PBS. Brains were extracted from the skull and incubated in 4% paraformaldehyde in PBS overnight. Brains were suspended in 3% agarose in PBS and sectioned in ice-cold PBS with a vibratome (7000smz-2, Campden Instruments). 50 µm sections were collected and placed free-floating in well-plates filled with PBS. Every fourth section was stained with DAPI in PBS (1:2000) for one hour, washed 3×5 minutes in PBS, mounted on microscope slides (Superfrost Plus, Fisher Scientific), and coverslipped with Fluormount-G (Southern Biotech). Slides were imaged using an Olympus VS200 Slide Scanner with a 20X objective using an X-Cite XYLIS (Excelitas Technologies) broad spectrum LED light that was maintained at 100% power. Cerebellum injection sites and the axonal projections from both cerebellum and IC were imaged using an exposure time of 1.493 ms for DAPI (378/52 nm excitation filter, 432/36 nm emission filter), and 16.617 ms exposure time for GFP (474/27 nm excitation filter, 515/30 emission filter). Many images of the IC injection site were saturated with these exposure times, so IC was also imaged with a reduced exposure time of 2.53 ms for GFP. Additional 63X images were acquired using a Zeiss LSM 800 confocal microscope.

### Image analysis

To address variations in background fluorescence in the 20X slide scanner images, background subtraction and thresholding functions were used in FIJI (Schindelin et al., 2012). Regions of interest were made and watershed and analyze particle functions in FIJI were used to count the number of labeled somas within the IC. All labeled cells in the cerebellum were counted using the same functions including both cerebellar cortex and cerebellar nuclei. Labeled volume is a measure that represents a combination of labeling density and distance of spread and was calculated as the number of pixels above a brightness threshold of 20 across all collected slices. Each pixel was 0.325 µm^2^, the depth was the thickness of the slice (50 µm) and, the final volume was multiplied by four to account for imaging every fourth slice. Mean pixel brightness was calculated as the average of the pixel intensities that were above threshold in the image.

The ability of the AAV serotypes to label different cell types in the cerebellar cortex was analyzed in FIJI (Schindelin et al., 2012). Regions of interest were created using the wand tool to outline individual labeled cells and cell types were annotated and counted based on their morphology. Purkinje cells were identified by their large soma size, extensive dendritic tree, and location in the Purkinje cell layer.

Unipolar brush cells were identified by their location in the granule cell layer and unique brush dendrite. Golgi cells were identified by their large size and presence in the granule cell layer. Granule cells were determined by their small size, absence of dendritic brush, and location in the granule cell layer. The labeling intensity varied across granule cells from bright to quite dim across serotypes. Some labeled granule cells were not markedly brighter than the surrounding neuropil (Kim et al., 2015). However, the observation that many DAPI positive cells were completely devoid of any GFP fluorescence (Figure 3B, white arrows), justified identifying these dimly labeled cells as expressing the transgene. There was no apparent difference in the proportion of brightly labeled granule cells across serotypes. Molecular layer interneurons were determined by their small size and presence in the molecular layer. 9-14 images were analyzed for each serotype using 2-4 images (567.90 µm x 567.90 µm) per animal (4 animals for AAV1, 2, 5, 8, 9, and 3 animals for AAVrg) per serotype.

To analyze retrograde labeling for each serotype, the number of labeled cell bodies in each region was counted manually, using the Allen Institute mouse brain atlas as a reference. To analyze anterograde axonal projections, the background subtraction function in FIJI was applied and the total number of pixels above the threshold was used as an estimate of the density of labeled axons. One image per animal per region (3-4 per serotype) was analyzed for each brain region for both retrograde and anterograde labeling studies.

Identical imaging parameters and brightness and contrast adjustments were applied to all images within each figure panel that were included in quantitative comparisons. Non-linear adjustments (gamma) were not applied to any image.

### Statistics

The Shapiro-Wilk test was used to determine whether the data were normally distributed. Groups were compared using one-way ANOVAs followed by Tukey’s HSD tests. Statistical analyses are not reported for data where more than two of the serotypes failed the Shapiro-Wilk test and instead qualitative comparisons are reported. Prism (Graphpad) and R were used for statistical tests and for generating figures.

## Results

The aim of this study was to test the efficacy of different AAV serotypes in transducing neurons in the IC and cerebellum of mice. Identical titers (8*10^12^ GC/ml) and volumes (500 nl) of AAV-hSyn-GFP constructs in the AAV serotypes 1, 2, 5, 8, 9, and the directed evolution derived AAVrg (Tervo et al., 2016), were stereotaxically injected into the brains of anesthetized mice. The mice quickly recovered from the procedure and did not show any changes in behavior or health during the two-week incubation period. For each region, at least four animals were used for each serotype. Following transcardiac perfusion, brain extraction, and fixation, 50 µm serial sections were collected, and every fourth section was stained with DAPI, mounted on slides, imaged, and analyzed.

### Efficacy of transgene expression in the inferior colliculus

First, we tested the ability of the most widely available AAV serotypes to drive transgene expression in the IC. The AAV injections were made into the central IC and the volume of 500 nl was sufficient to diffuse to all regions including the dorsal and external nuclei. GFP was expressed by all serotypes, in all experiments (Figure 1A). In some figures, the serotype AAVrg is shown for comparison, but is not included in statistical analyses of transduction volume and brightness. AAV1 drove the highest GFP expression in the IC compared to the other serotypes with a substantial degree of spread (Figure 1A-B), while AAV5 had the lowest GFP expression (Figure 1A). To quantitatively compare the transduction efficacy across serotypes, we calculated the total number of labeled cells within the IC of the imaged sections. Significantly more cells were labeled by AAV1 than AAV2, 5, 8, and 9 (Figure 1C, Table 1). Additionally, AAV8 showed a significantly greater number of labeled cells than AAV2 and 5 (Figure 1C). AAV1 labeled more cells outside the IC in the adjacent periaqueductal gray than all other serotypes (One-way ANOVA, F (4, 15) = 16.44, P < 0.0001, Tukey’s HSD tests, AAV1 vs AAV2, 5, 8, P < 0.001; AAV1 vs AAV9, P = 0.0009). Although AAV9 labeled far fewer cells in IC than did AAV1, the percentage of cells labeled outside the IC of the total population was similar between them, indicating similar spread (AAV1: 10.6 ± 1.98 %, AAV9: 11.0 ± 2.78 %), while the other AAVs labeled few cells outside IC resulting in labeling that was more restricted to the injection site (AAV2: 3.6 ± 1.0 %, AAV5: 1.9 ± 0.4 %, AAV8: 1.6 ± 1.2 %; mean ± SEM). The volume of viral labeling was determined by summing the total number of GFP positive pixels across all slices. AAV1 transduced a significantly greater volume compared to AAV2, 5, 8, and 9 (Figure 1D). Finally, the mean pixel brightness for each serotype was calculated, showing cells infected with AAV1 had significantly brighter GFP expression compared to those infected with AAV2, 5, and 9 (Figure 1E). In sum, AAV1 labeled the most IC neurons and labeled the largest volume, while serotypes AAV2, 5, 8, and 9 were far less effective (Figure 1F).

**Figure 1:**
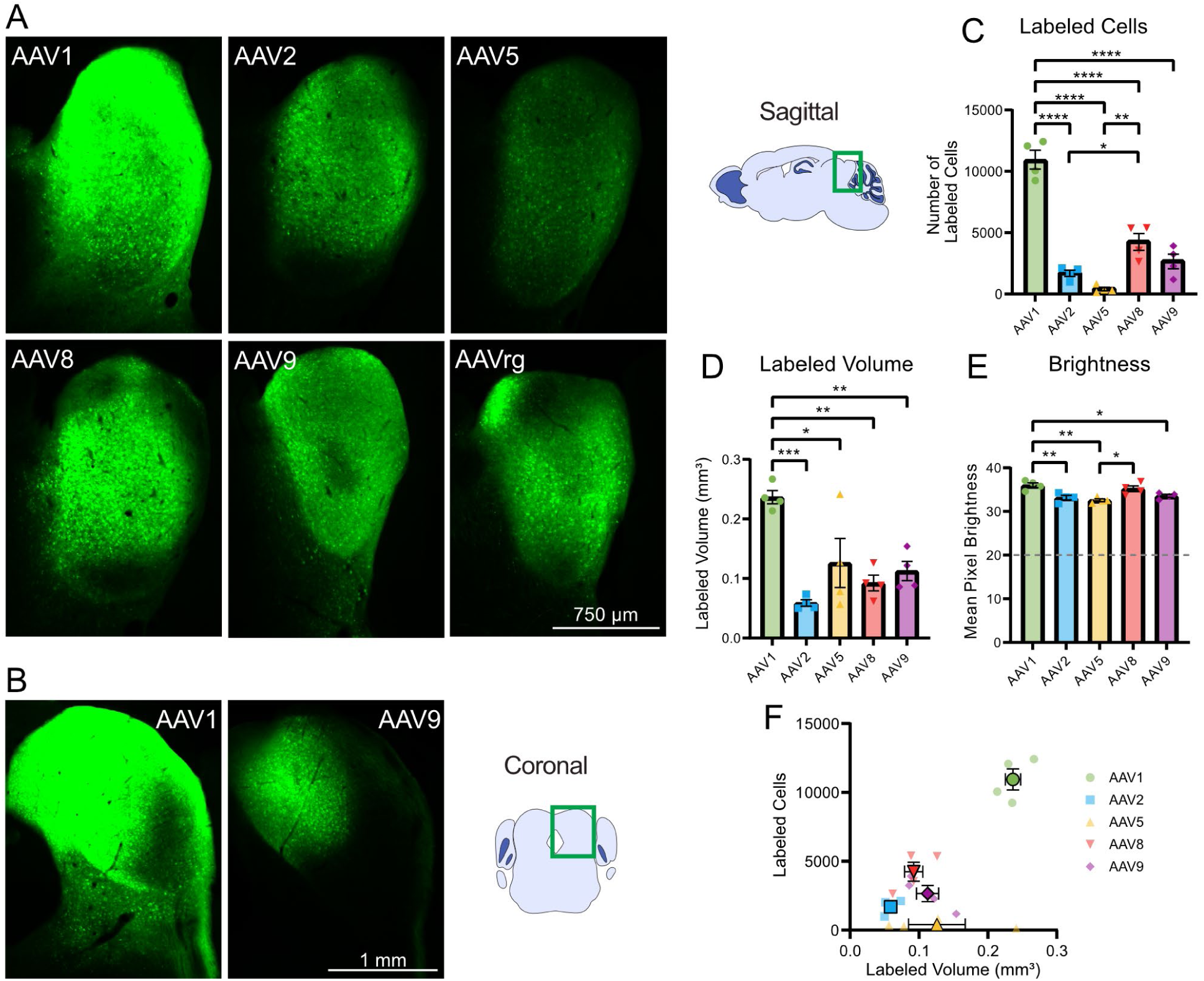
AAV1 produced the highest transgene expression and the largest labeled volume in the inferior colliculus. (A) Intracranial injection of the AAV1, 2, 5, 8, 9, rg serotypes led to GFP expression in the inferior colliculus. Representative sagittal sections of the injection site in the inferior colliculus for each serotype. (B) Additional representative coronal sections of AAV1 and AAV9 injected inferior colliculus. (C) Bar graph showing mean number of labeled cells across serotypes. AAV1 labeled significantly more cells than AAV2, 5, 8, and 9, indicating superior transduction efficacy. (D) Bar graph showing mean volume of labeling across serotypes. The volume of AAV1 labeling also was significantly greater than that of AAV 2, 5, 8, and 9. Horizontal line indicates the brightness threshold over which pixels were considered labeled and were included in the mean brightness calculation. (E) Bar graph showing mean pixel brightness across serotypes. AAV1 produced significantly greater mean pixel brightness than AAV 2, 5, and 9. (F) Comparison of number of labeled cells with labeled volume. Identical imaging parameters and brightness and contrast adjustments were applied to all images in the same panel. Statistical comparisons were made using one-way ANOVAs followed by Tukey’s HSD tests. Error bars are SEM. * indicates P<0.05, ** indicates P<0.01, *** indicates P<0.001, **** indicates P<0.0001.

**Table 1.** Statistics associated with Figure 1.

| Associated figure | Statistical test | Groups compared | Test statistic | P value | Mean difference | N (animals) |
| --- | --- | --- | --- | --- | --- | --- |
| Fig 1C | One-way ANOVA | All | $F(4, 15) = 56.76$ | <0.0001 | | 20 |
| Fig 1C | Tukey's test | AAV1 vs. AAV2 | $q = 16.87$ | <0.0001 | 9260 | 8 |
| Fig 1C | Tukey's test | AAV1 vs. AAV5 | $q = 19.21$ | <0.0001 | 10547 | 8 |
| Fig 1C | Tukey's test | AAV1 vs. AAV8 | $q = 12.22$ | <0.0001 | 6707 | 8 |
| Fig 1C | Tukey's test | AAV1 vs. AAV9 | $q = 15.11$ | <0.0001 | 8294 | 8 |
| Fig 1C | Tukey's test | AAV5 vs. AAV8 | $q = 6.993$ | 0.0014 | -3840 | 8 |
| Fig 1C | Tukey's test | AAV2 vs. AAV8 | $q = 4.650$ | 0.0343 | -3840 | 8 |
| Fig 1D | One-way ANOVA | All | $F(4, 15) = 9.842$ | 0.0004 | | 20 |
| Fig 1D | Tukey's test | AAV1 vs. AAV2 | $q = 8.301$ | 0.0003 | 0.1775 | 8 |
| Fig 1D | Tukey's test | AAV1 vs. AAV5 | $q = 5.161$ | 0.0172 | 0.1104 | 8 |
| Fig 1D | Tukey's test | AAV1 vs. AAV8 | $q = 6.736$ | 0.0020 | 0.1441 | 8 |
| Fig 1D | Tukey's test | AAV1 vs. AAV9 | $q = 5.792$ | 0.0072 | 0.1239 | 8 |
| Fig 1E | One-way ANOVA | All | $F(4, 15) = 8.588$ | 0.0008 | | 20 |
| Fig 1E | Tukey's test | AAV1 vs. AAV2 | $q = 5.759$ | 0.0076 | 2.836 | 8 |
| Fig 1E | Tukey's test | AAV1 vs. AAV5 | $q = 6.963$ | 0.0015 | 3.429 | 8 |
| Fig 1E | Tukey's test | AAV1 vs. AAV9 | $q = 4.943$ | 0.0231 | 2.434 | 8 |
| Fig 1E | Tukey's test | AAV5 vs. AAV8 | $q = 5.350$ | 0.0133 | -2.635 | 8 |

### Efficacy of transgene expression in the cerebellum

These AAVs were also tested in the cerebellum, where the injections were focused on the lateral aspect of lobule X. We chose this region because lobule X is part of the vesibulocerebellum and contains a high density of unipolar brush cells (UBCs), in addition to the other cell types present throughout all lobes of the cerebellar cortex. Additionally, the lateral injection site allowed for transfection of the cerebellar nuclei. AAV1 labeled significantly more cerebellar neurons than did AAV2, 5, 8, and 9 (Figure 2A-D, Table 2). The volume of brain labeled by AAV1 was also significantly greater than that of AAV2, 5, 8, and 9 (Figure 2E). Comparing the mean pixel brightness across serotypes revealed that AAV1 labeling was brighter than the other serotypes and reached significance compared to AAV5 (Figure 2F). Thus, as in the IC, AAV1 labeled the most neurons and labeled the largest volume while the other serotypes produced much more restricted expression (Figure 2G).

**Figure 2:**
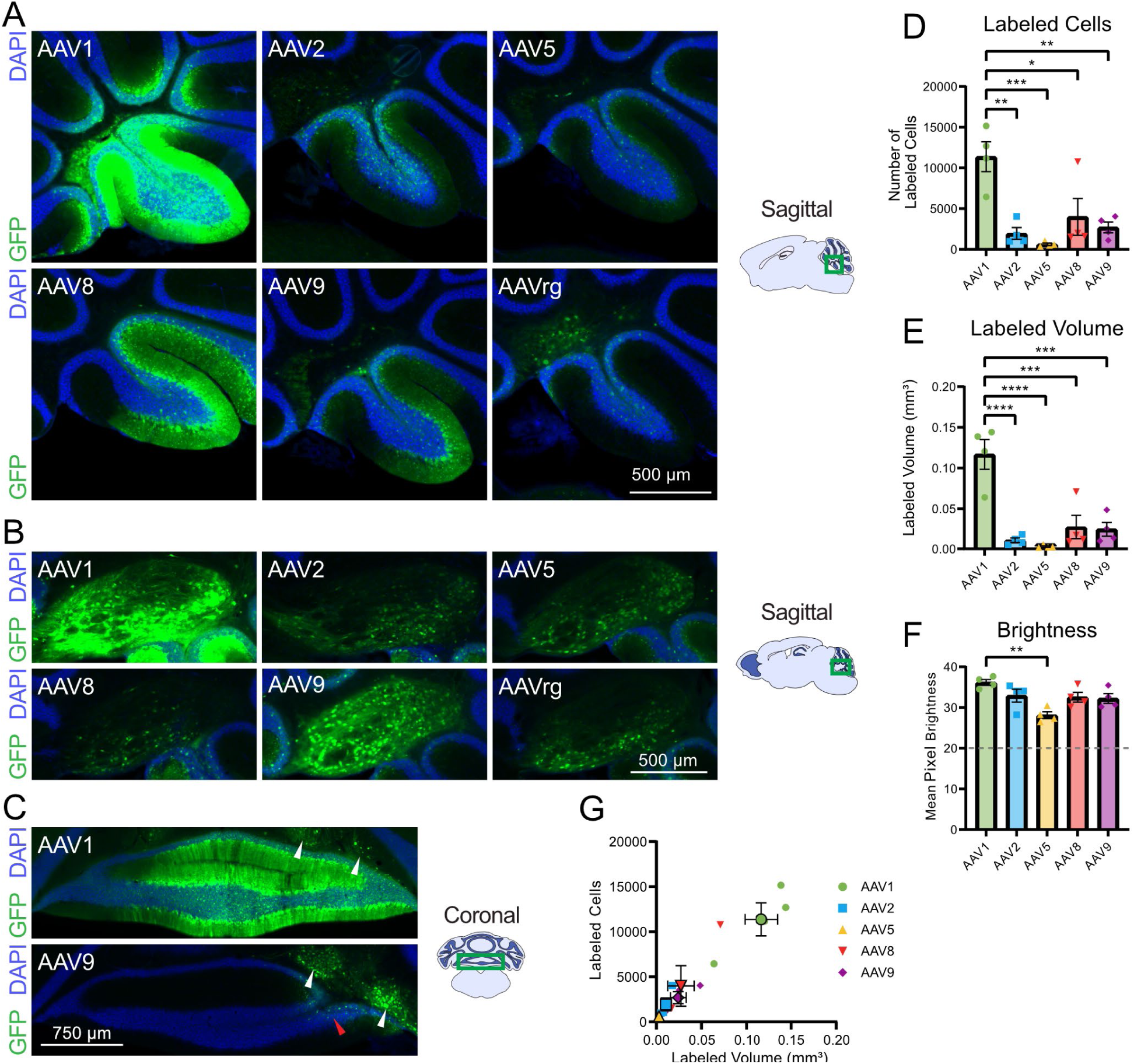
AAV1 produced the highest transgene expression and largest labeled volume in the cerebellum. (A) Representative sagittal sections of injected cerebellar cortex for each serotype. (B) Representative sagittal sections of the interposed nucleus. (C) Additional representative coronal sections of AAV1 and AAV9 injected cerebellum. White arrows indicate transfected cells in the deep cerebellar nuclei, the red arrow indicates the minimal spread to the cerebellar cortex by AAV9. (D) Bar graph showing mean number of labeled cells across serotypes. AAV1 labeled significantly more cells than AAV2, 5, 8, and 9, indicating superior transduction efficacy. (E) Bar graph showing mean volume of labeling across serotypes. The labeled volume of AAV1 also was significantly greater than that of AAV 2, 5, 8, and 9. (F) Bar graph showing mean pixel brightness across serotypes. AAV1 produced significantly greater mean pixel brightness compared to AAV 2, 5, 8, and 9. Horizontal line indicates the brightness threshold over which pixels were considered labeled and were included in the mean brightness calculation. (G) Comparison of number of cells labeled by labeled volume. Identical imaging parameters and brightness and contrast adjustments were applied to all images in the same panel. Statistical comparisons were made using one-way ANOVAs followed by Tukey’s HSD tests. Error bars are SEM. * indicates P<0.05, ** indicates P<0.01, *** indicates P<0.001, **** indicates P<0.0001.

**Table 2.** Statistics associated with Figure 2.

| Associated figure | Statistical test | Groups compared | Test statistic | P value | Mean difference | N (animals) |
| --- | --- | --- | --- | --- | --- | --- |
| Fig 2D | One-way ANOVA | All | $F(4, 15) = 9.440$ | 0.0005 | | 20 |
| Fig 2D | Tukey's test | AAV1 vs. AAV2 | $q = 6.843$ | 0.0017 | 9418 | 8 |
| Fig 2D | Tukey's test | AAV1 vs. AAV5 | $q = 7.808$ | 0.0005 | 10746 | 8 |
| Fig 2D | Tukey's test | AAV1 vs. AAV8 | $q = 5.366$ | 0.0130 | 7384 | 8 |
| Fig 2D | Tukey's test | AAV1 vs. AAV9 | $q = 6.300$ | 0.0036 | 8671 | 8 |
| Fig 2E | One-way ANOVA | All | $F(4, 15) = 16.65$ | <0.0001 | | 20 |
| Fig 2E | Tukey's test | AAV1 vs. AAV2 | $q = 9.408$ | <0.0001 | 0.1062 | 8 |
| Fig 2E | Tukey's test | AAV1 vs. AAV5 | $q = 10.09$ | <0.0001 | 0.1139 | 8 |
| Fig 2E | Tukey's test | AAV1 vs. AAV8 | q = 7.938 | 0.0004 | 0.08958 | 8 |
| Fig 2E | Tukey's test | AAV1 vs. AAV9 | q = 8.199 | 0.0003 | 0.09252 | 8 |
| Fig 2F | One-way ANOVA | All | F (4, 15) = 6.205 | 0.0037 |  | 20 |
| Fig 2F | Tukey's test | AAV1 vs. AAV5 | q = 6.999 | <0.0001 | 8.064 | 8 |

### Analysis of cell-type tropism for AAV serotypes in the cerebellum

The cerebellum is highly organized with three distinct cortical layers–the granule cell layer, Purkinje cell layer, and molecular layer–and has a host of cell types that can be identified by their location within these layers and by their unique morphologies (see Methods). The synapsin promoter ensures transgene expression is only driven in neurons (Kügler et al., 2003), thus allowing for visual analysis of cell-type tropism within the cerebellum. High-resolution images of DAPI and GFP in lobule X of the cerebellum were imaged on a confocal microscope (Figure 3A). Fluorescently labeled cells were confirmed to contain a DAPI stained nucleus, and were manually annotated by cell-type (Figure 3B).

**Figure 3:**
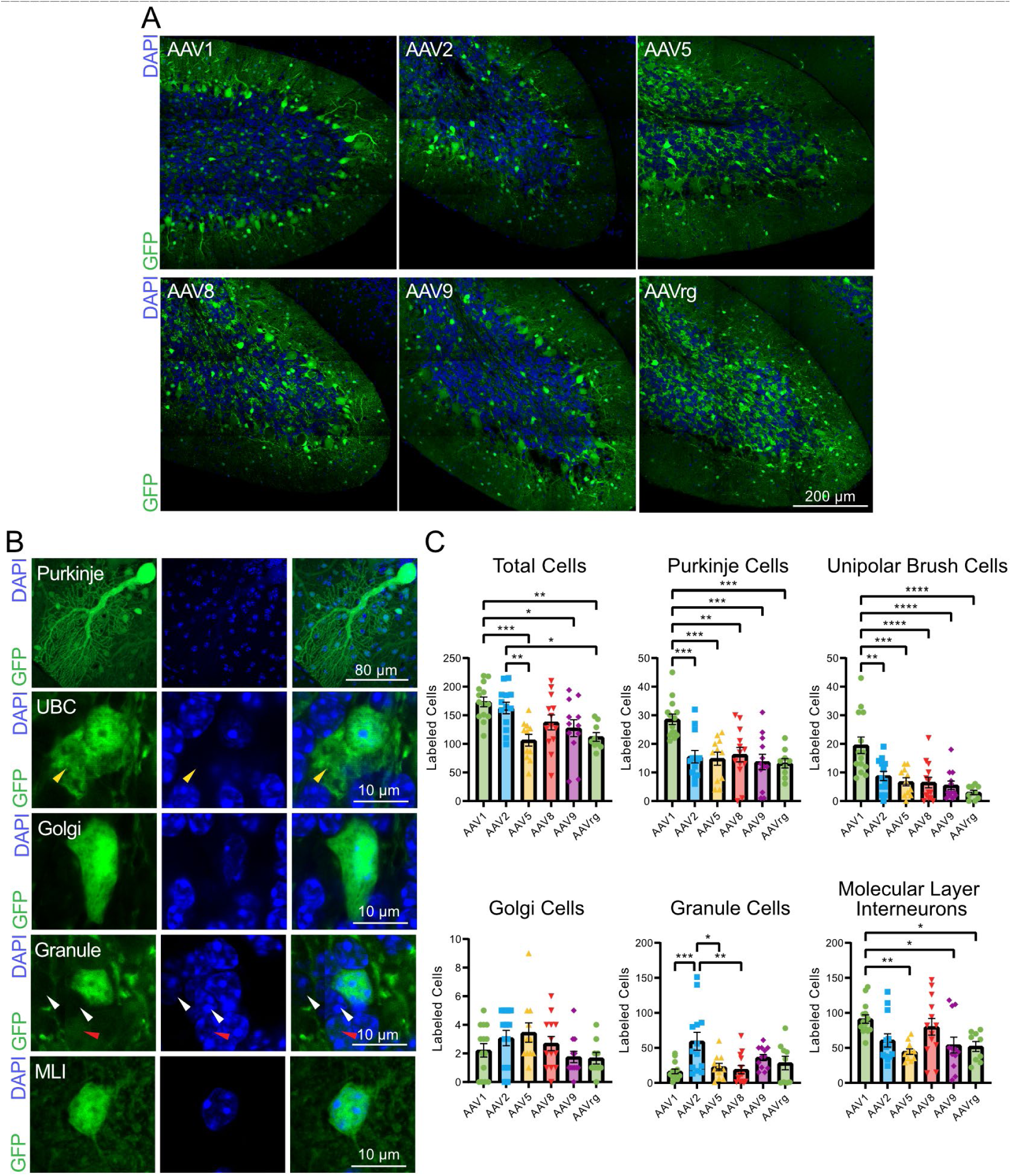
AAV serotype tropism in the cerebellum. (A) Representative sagittal sections of lobule X of the cerebellum. (B) Representative images of cerebellar cell types. Fluorescent cells were characterized utilizing the unique morphology of cerebellar cortex cell types. The yellow arrow indicates the dendritic brush unique to unipolar brush cells. White arrows indicate DAPI positive cells devoid of GFP expression and the red arrow indicates a DAPI positive cell with dim GFP expression. (C) Bar graphs showing the number of GFP positive cerebellar cell types across AAV serotypes. 63X images of lobule X of the cerebellum were analyzed to determine tropism of AAV 1, 2, 5, 8, 9, and rg. The AAV1 serotype labeled significantly more total cells compared to AAV5, 9, and rg. AAV1 labeled significantly more Purkinje and unipolar brush cells compared to all the other serotypes, and significantly more molecular layer interneurons than AAV5, 9, and rg. AAV2 transfected significantly more granule cells compared to AAV1, 5, and 8. Identical imaging parameters and brightness and contrast adjustments were applied to all images in the same panel. Statistical comparisons were made using one-way ANOVAs followed by Tukey’s HSD tests. Error bars are SEM. * indicates P<0.05, ** indicates P<0.01, *** indicates P<0.001, **** indicates P<0.0001.

This analysis revealed that serotypes varied in their ability to transfect different cerebellar cell types. As expected, AAV1 labeled significantly more total cells, although AAV2 also labeled many cells, despite the relatively low labeled volume shown in the previous figure (Figure 3C, Table 3). AAV1 was the most effective across cell types, and labeled significantly more Purkinje and unipolar brush cells compared to all the other serotypes, and significantly more molecular layer interneurons compared to AAV5, 9, and rg (Figure 3C). However, AAV2 transfected significantly more granule cells than the other serotypes and reached significance when compared to AAV1, 5, and 8 (Figure 3C). Thus, AAV1 labeled the largest variety of cell types, whereas AAV2 labeled granule cells most effectively.

**Table 3.** Statistics associated with Figure 3.

| Associated figure | Statistical test | Groups compared | Test statistic | P value | Mean difference | N (slices) |
| --- | --- | --- | --- | --- | --- | --- |
| Fig 3C- Total cells | One-way ANOVA | All | F (6, 72) = 5.771 | 0.0002 |  | 72 |
| Fig 3C- Total cells | Tukey's test | AAV1 vs. AAV5 | q = 6.042 | 0.0009 | 66.99 | 25 |
| Fig 3C- Total cells | Tukey's test | AAV1 vs. AAV9 | q = 4.243 | 0.0423 | 45.94 | 26 |
| Fig 3C- Total cells | Tukey's test | AAV1 vs. AAVrg | q = 5.209 | 0.0060 | 61.25 | 23 |
| Fig 3C- Total cells | Tukey's test | AAV2 vs. AAV5 | q = 5.003 | 0.0093 | 56.41 | 24 |
| Fig 3C- Total cells | Tukey's test | AAV2 vs. AAVrg | q = 4.245 | 0.0421 | 50.66 | 22 |
| Fig 3C- Purkinje cells | One-way ANOVA | All | F (6, 72) = 7.164 | <0.0001 |  | 72 |
| Fig 3C- Purkinje cells | Tukey's test | AAV1 vs. AAV2 | q = 6.096 | 0.0008 | 13.03 | 27 |
| Fig 3C-Purkinje<br>cells | Tukey's<br>test | AAV1 vs. AAV5 | q = 6.150 | 0.0007 | 13.75 | 25 |
| Fig 3C-Purkinje<br>cells | Tukey's<br>test | AAV1 vs. AAV8 | q = 5.772 | 0.0017 | 12.34 | 27 |
| Fig 3C-Purkinje<br>cells | Tukey's<br>test | AAV1 vs. AAV9 | q = 6.826 | 0.0001 | 14.90 | 26 |
| Fig 3C-Purkinje<br>cells | Tukey's<br>test | AAV1 vs. AAVrg | q = 6.472 | 0.0003 | 15.35 | 23 |
| Fig 3C-UBCs | One-way<br>ANOVA | All | F (6,<br>72) = 9.435 | <0.0001 |  | 72 |
| Fig 3C-UBCs | Tukey's<br>test | AAV1 vs. AAV2 | q = 5.830 | 0.0014 | 10.73 | 27 |
| Fig 3C-UBCs | Tukey's<br>test | AAV1 vs. AAV5 | q = 6.634 | 0.0002 | 12.77 | 25 |
| Fig 3C-UBCs | Tukey's<br>test | AAV1 vs. AAV8 | q = 7.084 | <0.0001 | 13.04 | 27 |
| Fig 3C-UBCs | Tukey's<br>test | AAV1 vs. AAV9 | q = 7.447 | <0.0001 | 14.00 | 26 |
| Fig 3C-UBCs | Tukey's<br>test | AAV1 vs. AAVrg | q = 8.082 | <0.0001 | 16.50 | 23 |
| Fig 3C-Golgi cells | One-way<br>ANOVA | All | F (6, 72) =<br>1.847 | 0.1157 |  | 72 |
| Fig 3C-Granule<br>cells | One-way<br>ANOVA | All | F (6, 72) =<br>4.792 | 0.0009 |  | 72 |
| Fig 3C-Granule cells | Tukey's test | AAV1 vs. AAV2 | q = 6.016 | 0.0009 | -43.03 | 27 |
| Fig 3C-Granule cells | Tukey's test | AAV2 vs. AAV5 | q = 4.828 | 0.0134 | 36.73 | 24 |
| Fig 3C-Granule cells | Tukey's test | AAV2 vs. AAV8 | q = 5.575 | 0.0026 | 40.62 | 26 |
| Fig 3C-MLIs | One-way ANOVA | All | F (6, 72) = 4.191 | 0.0023 |  | 72 |
| Fig 3C-MLIs | Tukey's test | AAV1 vs. AAV5 | q = 46.49 | 0.0055 | 5.243 | 25 |
| Fig 3C-MLIs | Tukey's test | AAV1 vs. AAV9 | q = 37.21 | 0.0381 | 4.299 | 26 |
| Fig 3C-MLIs | Tukey's test | AAV1 vs. AAVrg | q = 39.21 | 0.0482 | 4.171 | 25 |

### Cerebellar axonal projections

Expression of the AAV-hSyn-GFP transgene leads to cytosolic GFP expression and labeling throughout the entire neuron, including dendrites and axonal projections. Labeling axonal projections is useful to define projection targets of an injected area and is additionally significant to researchers utilizing AAVs to deliver genes encoding axonal proteins. All AAV serotypes labeled axonal projections to various regions distant from the injection site. These projections are evaluated for differences between serotypes below.

The thalamus receives axonal input from the cerebellum which is processed and conveyed to the cerebral cortex (Habas et al., 2019). The main targeted region is the ventral posterolateral nucleus of the thalamus (VPL) (Asanuma et al., 1983). Comparisons of these observations were made by summing the total number of labeled pixels per region, for each injection, per serotype. Projections to VPL were labeled following injection of AAV1 better than the other serotypes (Figure 4, Table 4). The red nucleus receives axonal input from the interposed nucleus of the cerebellar nuclei, which has been implicated in skilled reach and locomotor movement (Low et al., 2018; Judd et al., 2021). AAV1 and 9 produced more axonal labeling than did the other serotypes (Figure 4). These differences could reflect tropism of AAV1 and AAV9 to the interposed nucleus of the cerebellar nuclei.

**Figure 4:**
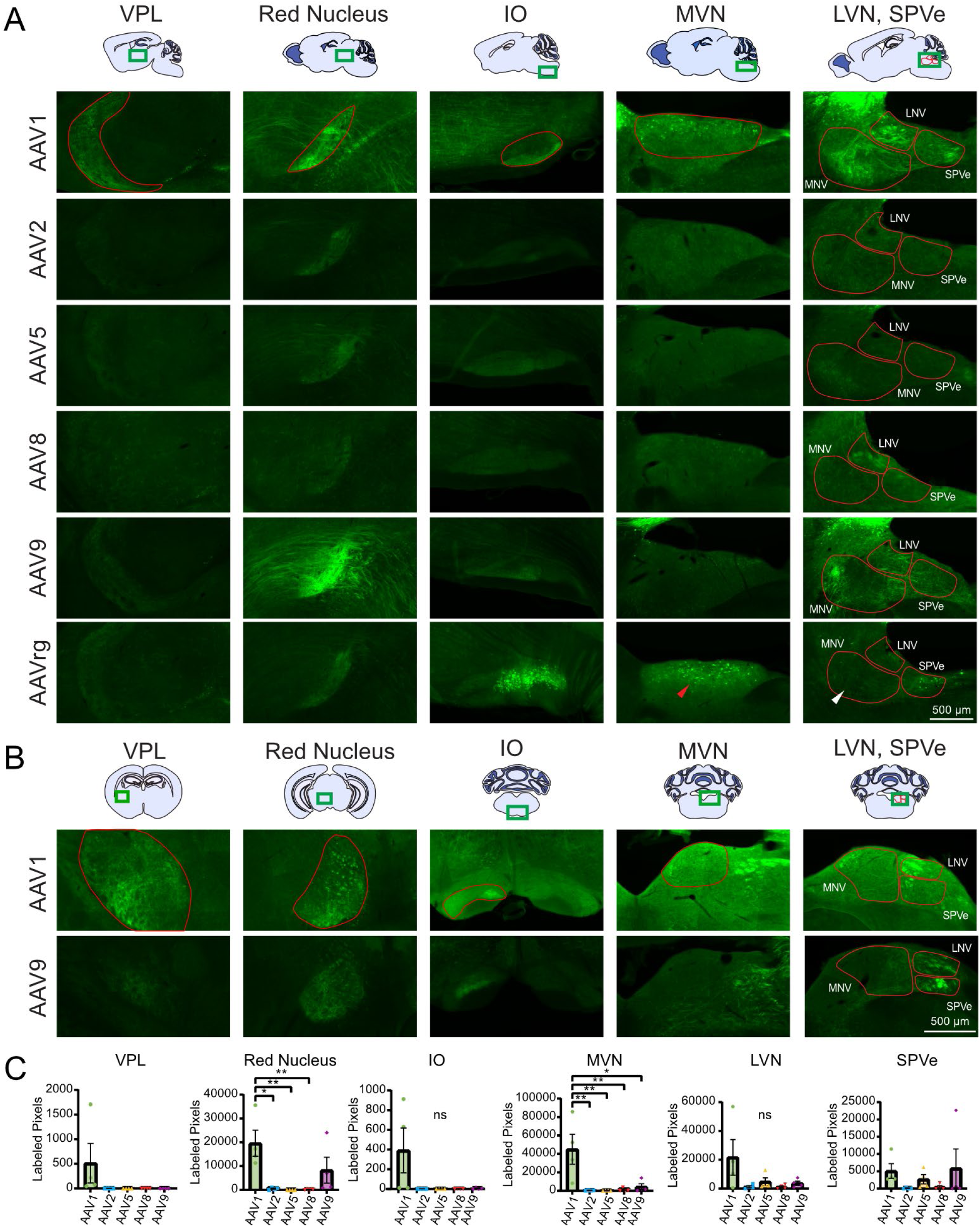
Labeled axonal projections from the cerebellum to the ventral posterolateral nucleus of the thalamus, the red nucleus, the inferior olive, the vestibular nuclei. (A) Representative images of the VPL, RN, IO, MVN, SPVe, and LVN by serotype. Differences in the degree of retrograde labeling within the MVN were present for slices infected with AAVrg (red arrow vs white arrow). (B) Representative coronal images of contralateral VPL, RN, IO, and the ipsilateral MVN, SPVe, and LVN. (C) Bar graphs showing serotype brightness quantification for each region. Labeled projections in all regions could be seen in slices infected AAV1 and AAV9, projections in the contralateral IO, ipsilateral MVN, LVN, and SPVe could be seen in slices infected with AAVrg, and projections in the MVN, LVN, and SPVe could be seen with slices infected with AAV8. Statistical comparisons were made using one-way ANOVAs followed by Tukey’s HSD tests. VPL and SPVe were not statistically tested. Error bars are SEM. * indicates P<0.05, ** indicates P<0.01, ns indicates P>0.05.

**Table 4.**
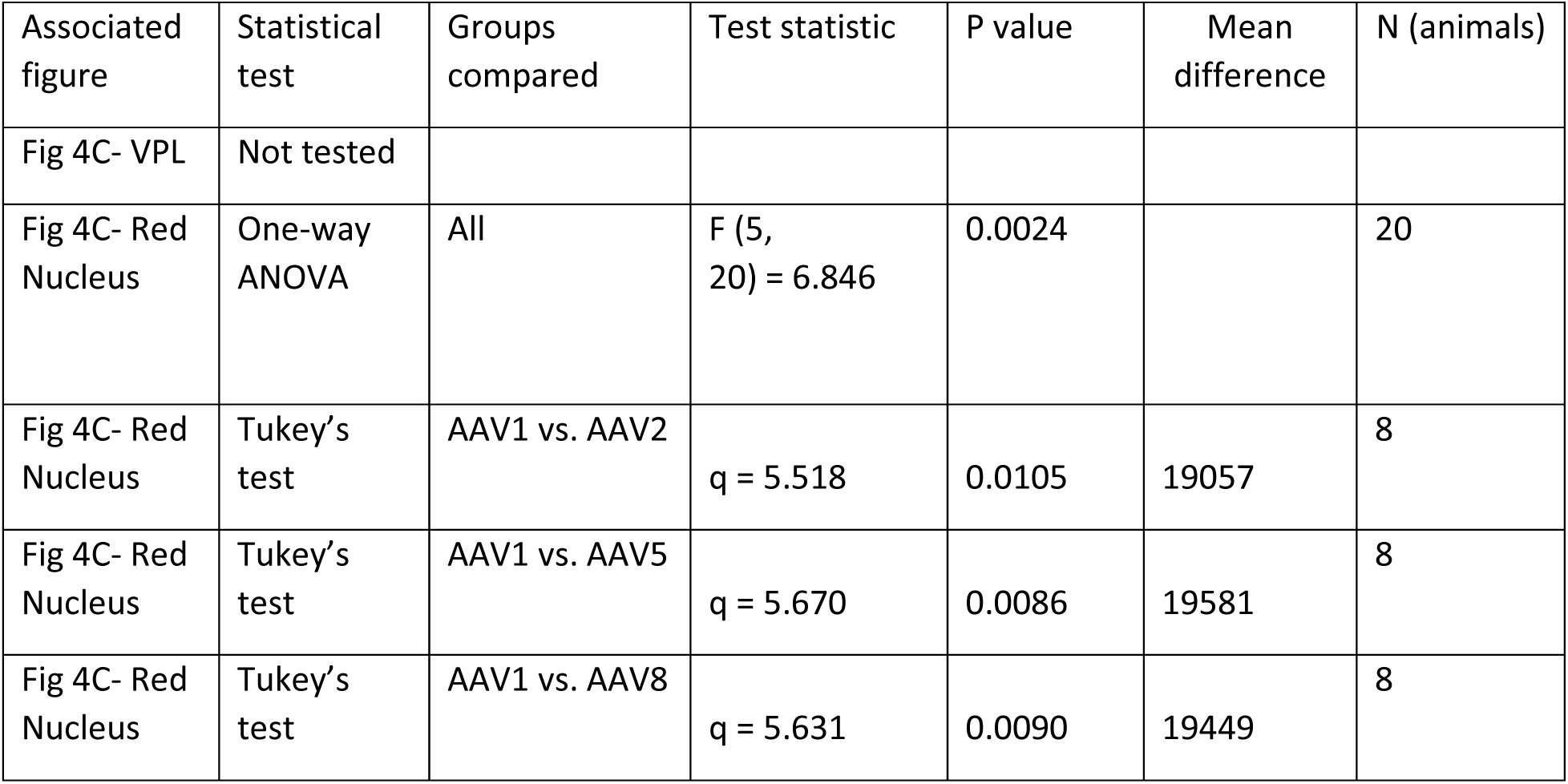
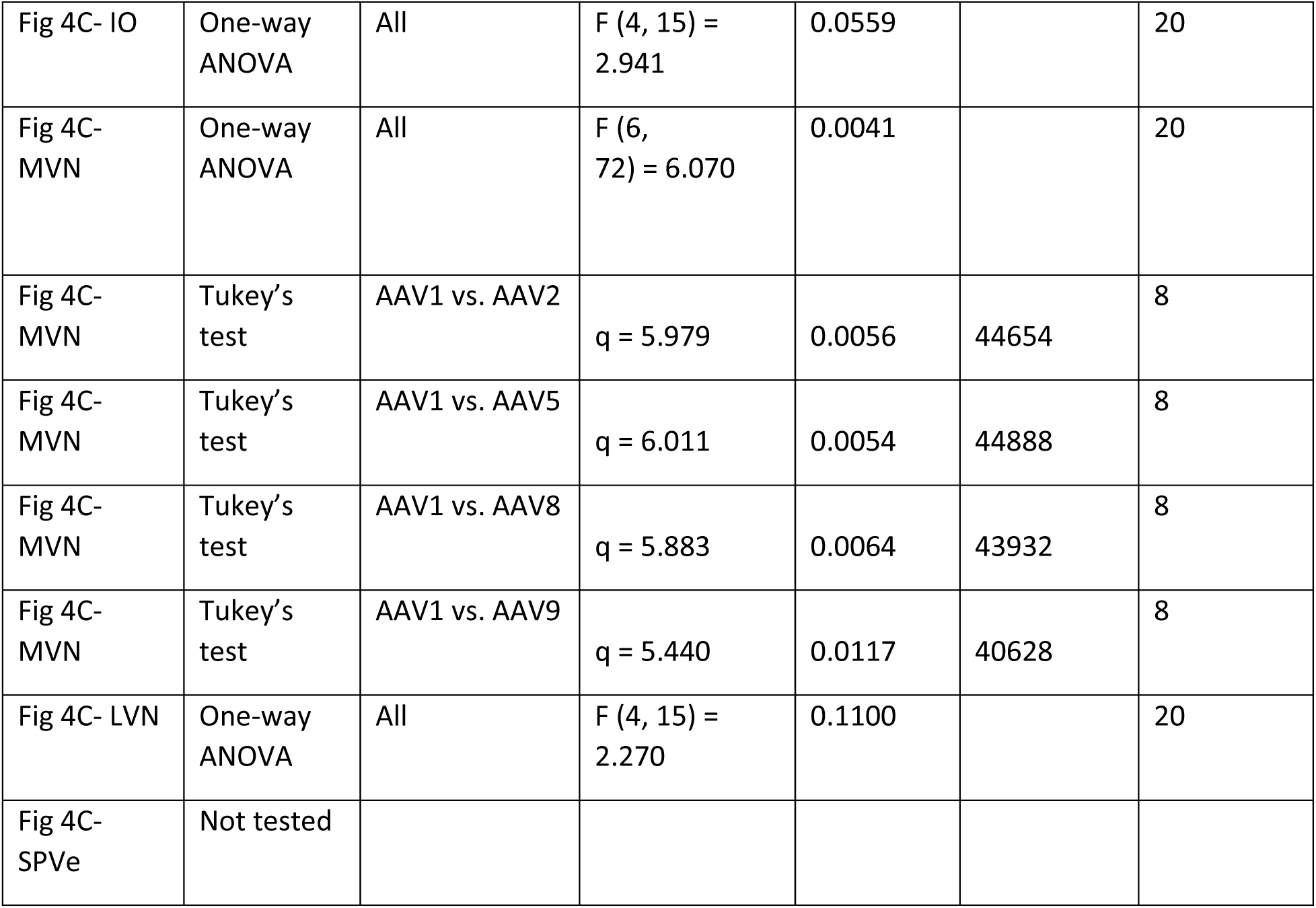
Statistics associated with Figure 4.

The inferior olive (IO) is the source of climbing fiber input to the cerebellum and is involved in learning and timing of movements, and comparing intended with achieved movements (De Zeeuw et al., 1998). The IO also receives descending input from the cerebellar nuclei. Labeled axonal projections were found in the contralateral inferior olive following injection of serotypes AAV1 (Figure 4).

The medial vestibular nucleus (MVN), lateral vestibular nucleus (LVN), and the spinal vestibular nucleus (SPVe) are three of the five vestibular nuclei where primary vestibular information is collected, processed, and modified by other sensory inputs (Barmack, 2023). Labeling was seen in the ipsilateral MVN, LVN, and SPVe following injection of AAV1 into the cerebellum (Figure 4). In the MVN, AAV1 demonstrated significantly more labeling than AAV2, 5, 8, and 9 (Figure 4). AAV1 had particularly greater labeling in the SPVe compared to AAV2, and 8.

Retrograde labeling was also seen in the ipsilateral vestibular nuclei where 2/8, 1/8, and 7/8 slices contained retrogradely labeled cells by AAV1, 5, and rg, respectively. The mean number of retrogradely labeled cells by AAVrg varied across the mediolateral extent of MVN, as seen in Figure 4A (white vs red arrows). The more medial regions of MVN in slices that did not contain LVN and SPVe had many more labeled cells following injection of AAVrg into the cerebellum (Figure 4A white arrow, mean ± SEM: 51.2 ± 9.3 labeled cells, n = 4) compared to more lateral regions of MVN containing all three vestibular nuclei (Figure 4A red arrow, mean ± SEM: 0.75 ± 0.25 labeled cells, n = 4). The observation that different zones of lobule X project to different vestibular nuclei may explain this finding (Wylie et al., 1994). Labeled cells were also observed in the LVN for 2/4 and 1/4 experiments by AAV1 and rg, respectively, while labeled cells in the SPVe were found in 1/4 and 4/4 experiments for AAV8 and rg, respectively. Additional retrograde labeling in other regions is explored below.

### Inferior collicular axonal projections

The DCN is a major source of ascending auditory input to the IC and the IC sends a large descending projection back to the DCN (Saldaña, 1993; Milinkeviciute et al., 2017; Balmer and Trussell, 2022). Labeled descending axonal projections were found in the ipsilateral DCN following injection of AAV1 to the IC with little-to-no labeling found following injection of AAV2, 5, 8, and 9, however AAV1 was only significantly greater than AAV5 with this small sample size (Figure 5, Table 5). Labeling in the contralateral DCN was seen following injection of AAVrg to the IC, and is likely a combination of axonal projections originating from labeled cells in the IC, DCN neurons that project to IC that are retrogradely labeled, and projections originating from retrogradely labeled cells in other brain regions. Meanwhile, sparse axonal labeling within in the contralateral DCN was seen following injection of AAV1 into the IC. Retrograde labeling was seen in the ipsilateral DCN for 2/4 AAVrg experiments, while labeling in the contralateral DCN was seen in 3/3 and 3/4 experiments for AAV1 and AAVrg, respectively.

**Figure 5:**
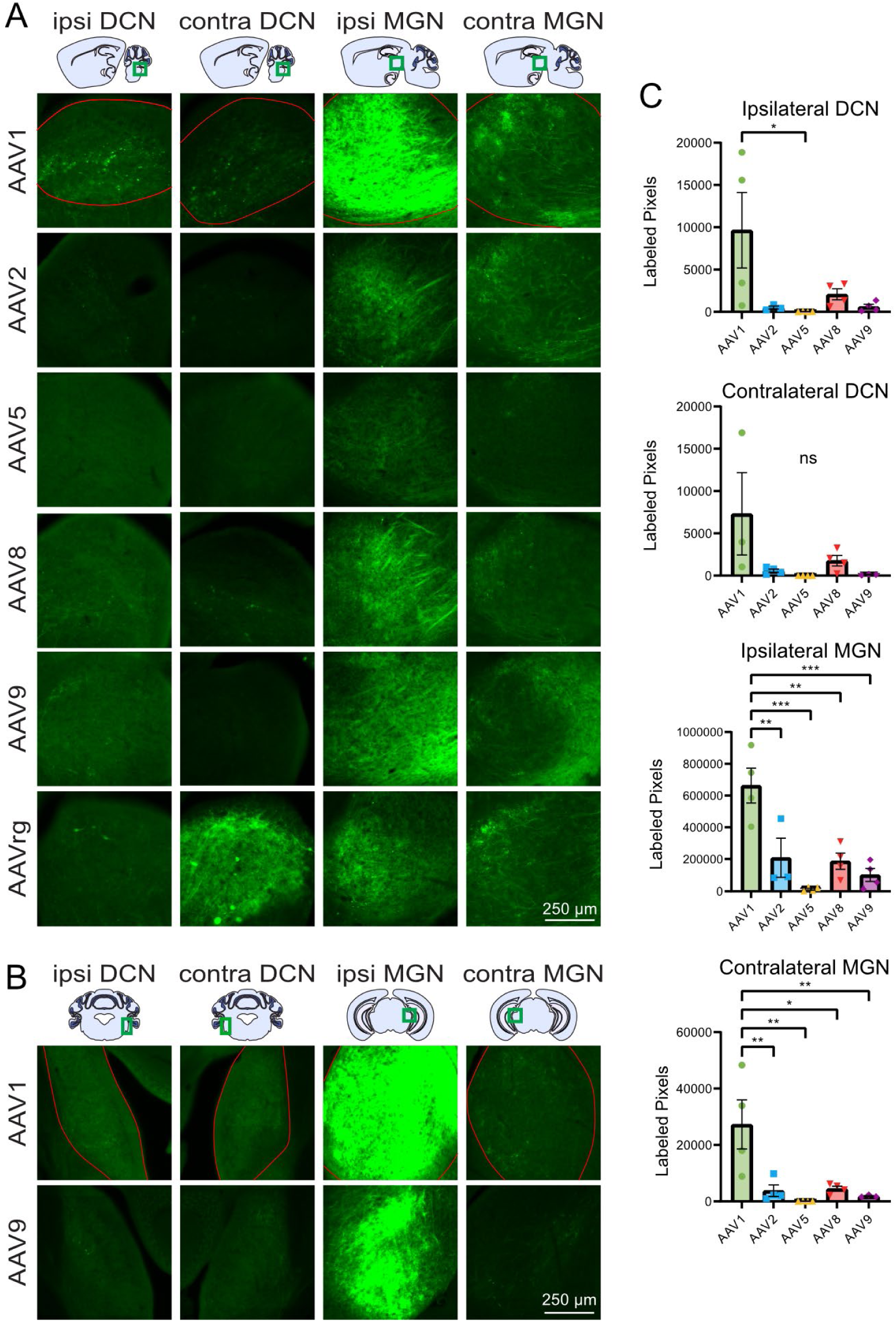
Labeled IC projections to the dorsal cochlear nucleus and medial geniculate nucleus. (A) Representative images of the DCN and MGN both ipsilateral and contralateral to the injection site. (B) Additional coronal images of the ipsilateral and contralateral DCN and MGN. (C) Bar graphs showing the mean serotype brightness quantification for each region. Statistical comparisons were made using one-way ANOVAs followed by Tukey’s HSD tests. Error bars are SEM. * indicates P<0.05, ** indicates P<0.01, *** indicates P<0.001, ns indicates P>0.05.

**Table 5.**
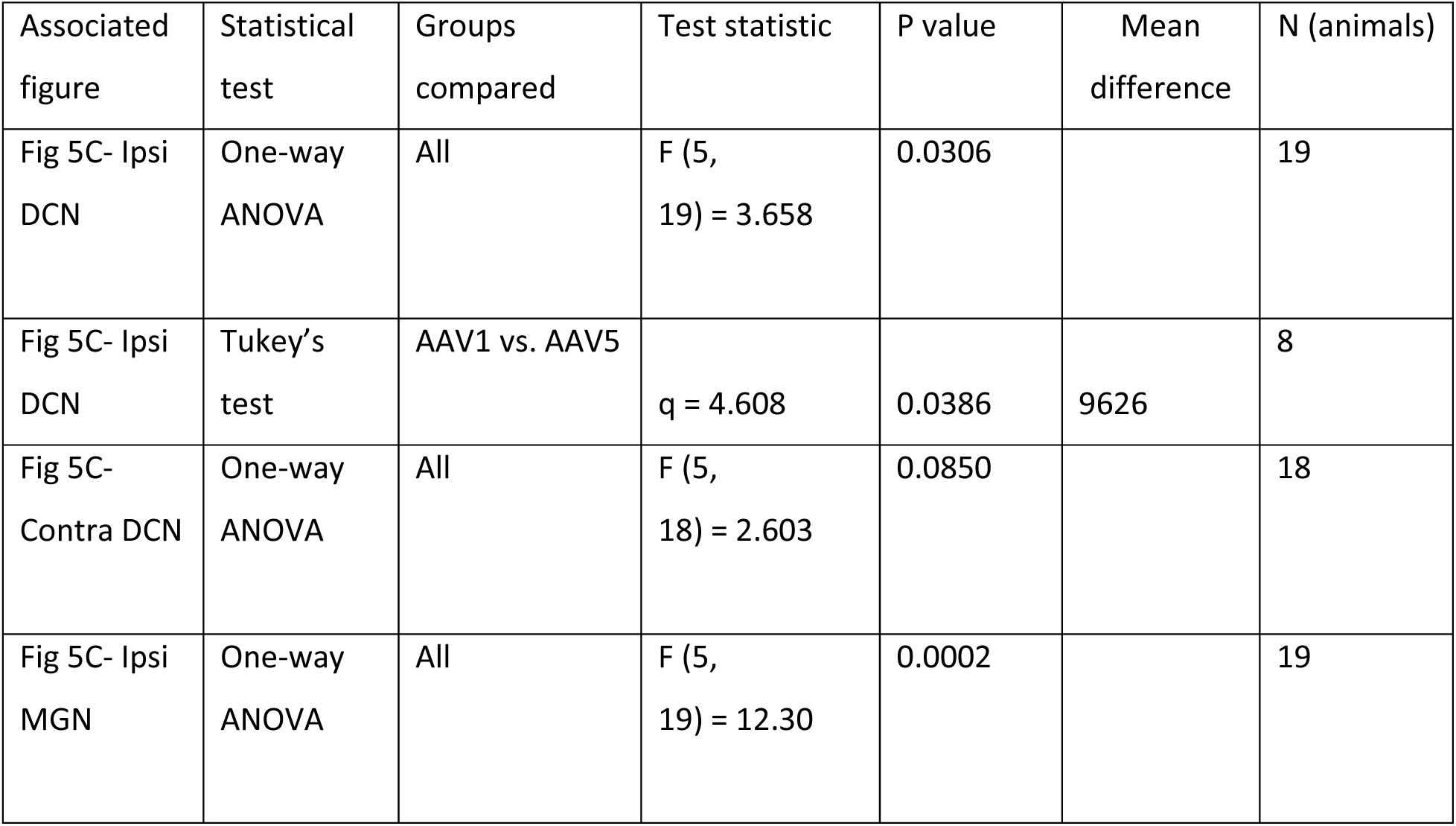
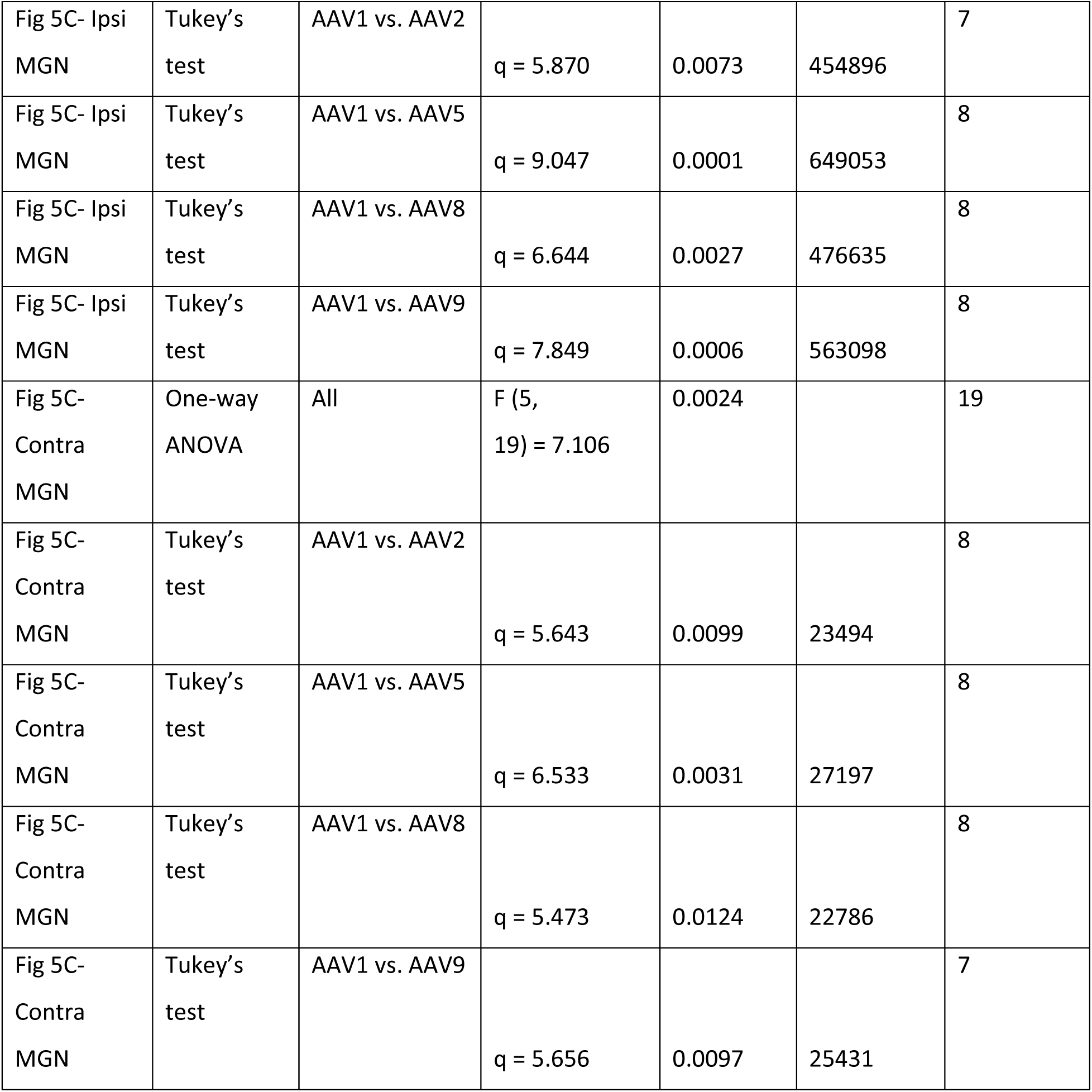
Statistics associated with Figure 5.

The main target of the IC is the ipsilateral MGN, which also integrates input from multiple ascending and descending sources (Winer and Schreiner, 2005). Extensive labeling was seen in both the ventral and dorsal divisions of the ipsilateral MGN for all serotypes with brightness decreasing in the following order: AAV1 >>> AAV2 > AAV8 > AAV9 > AAV5. AAV1 produced significantly brighter labeling in the ipsilateral MGN than all other serotypes (Figure 5B). Less intense labeling was seen in the contralateral MGN compared to the ipsilateral MGN following injection of the all the serotypes, however, AAV1 produced significantly greater labeling than AAV2, 5, 8, and 9 (Figure 5B). Across all experiments only one retrogradely labeled cell was found in the ipsilateral MGN following injection of AAVrg into the IC.

### Retrograde labeling of the nucleus of the lateral lemniscus by AAVs injected into the inferior colliculus

The lateral lemniscus is a fiber bundle that connects the cochlear nuclei and superior olivary complex with the IC (Felmy, 2019). The nucleus of the lateral lemniscus (NLL) is known for its generation of long-lasting inhibition in its contralateral counterpart and the IC (Felmy, 2019). Labeled cell bodies were found in the ipsilateral NLL following injections of AAV1, 5 and rg, indicating retrograde labeling of these cells that project to the IC. As expected, AAVrg retrogradely labeled the greatest number of NLL neurons (Figure 6). AAV2, 8, and 9 did not retrogradely label any cells in any experiments (Figure 6). However, AAV1 and AAV5 retrogradely labeled a number of neurons comparable to AAVrg (Figure 6).

**Figure 6:**
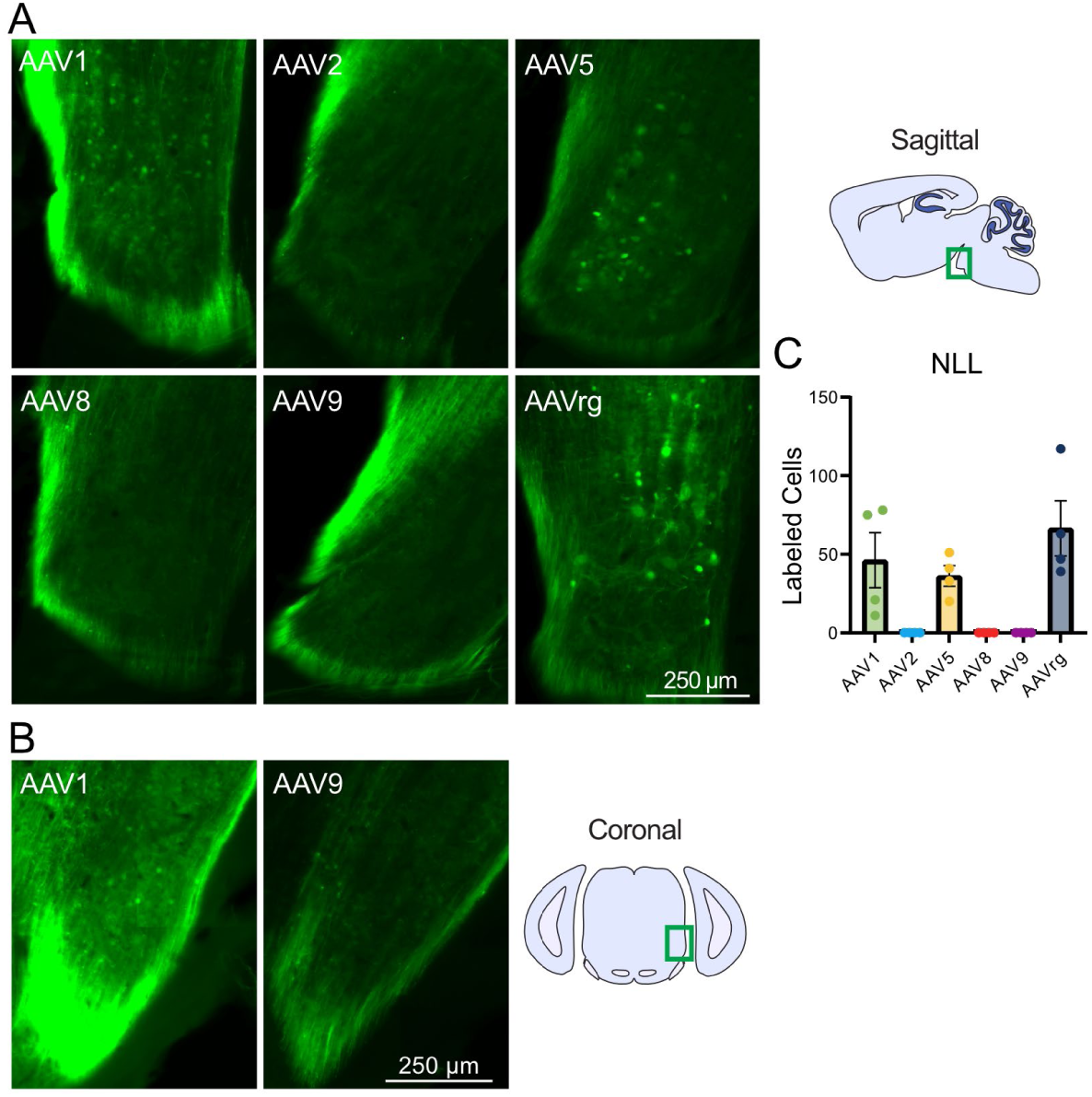
Retrograde labeling of the nucleus of the lateral lemniscus from the inferior colliculus. (A) Representative sagittal images of the ipsilateral nucleus of the lateral lemniscus by serotype. (B) Representative coronal images of the nucleus of the lateral lemniscus following AAV1 and 9 injection to the IC. (C) Bar graph showing mean number of labeled cells by serotype. Labeled cell bodies were present in the nucleus of the lateral lemniscus in brains injected with AAV1, 5, and rg, but completely absent AAV2, 8 or 9. Identical imaging parameters and brightness and contrast adjustments were applied to all images in the same panel. No statistical tests were performed.

### Retrograde labeling of the cuneate nucleus by AAVs injected into the cerebellum

The cuneate nucleus is a division of the dorsal column nuclei and receives primary sensory afferents from the upper body and upper limbs (Watson, 2012). AAV5 and AAVrg retrogradely labeled cell bodies in the ipsilateral cuneate nucleus (Figure 7). Labeled cell bodies were only rarely found in cuneate nucleus following cerebellar injections of other serotypes.

**Figure 7:**
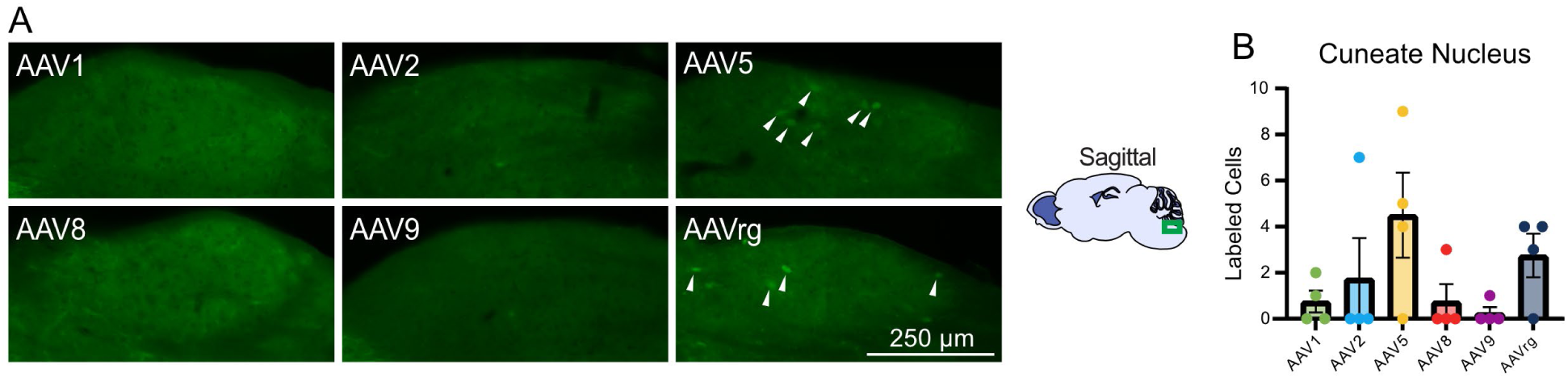
Retrograde labeling of the cuneate nucleus from the cerebellum. (A) Representative images of the ipsilateral cuneate nucleus by serotype. (B) Bar graph showing mean number of labeled cells by serotype. Labeled cell bodies were present in the cuneate nucleus in brains injected with AAV5, or rg (white arrows). Identical imaging parameters and brightness and contrast adjustments were applied to all images in the same panel. No statistical tests were performed.

### Summary of labeled axonal projections and retrograde labeling

The degree of retrograde labeling and presence of labeled axonal projections varied between serotypes and brain regions and are summarized in Figure 8. AAV1 labeled the most axonal projections in every area investigated (Figure 8A). All serotypes except AAV9 demonstrated some degree of retrograde labeling in at least one area, with AAVrg having the greatest degree of retrograde labeling, as expected (Figure 8B). The region with the greatest retrograde labeling across multiple serotypes was the nucleus of the lateral lemniscus (Figure 8B).

**Figure 8:**
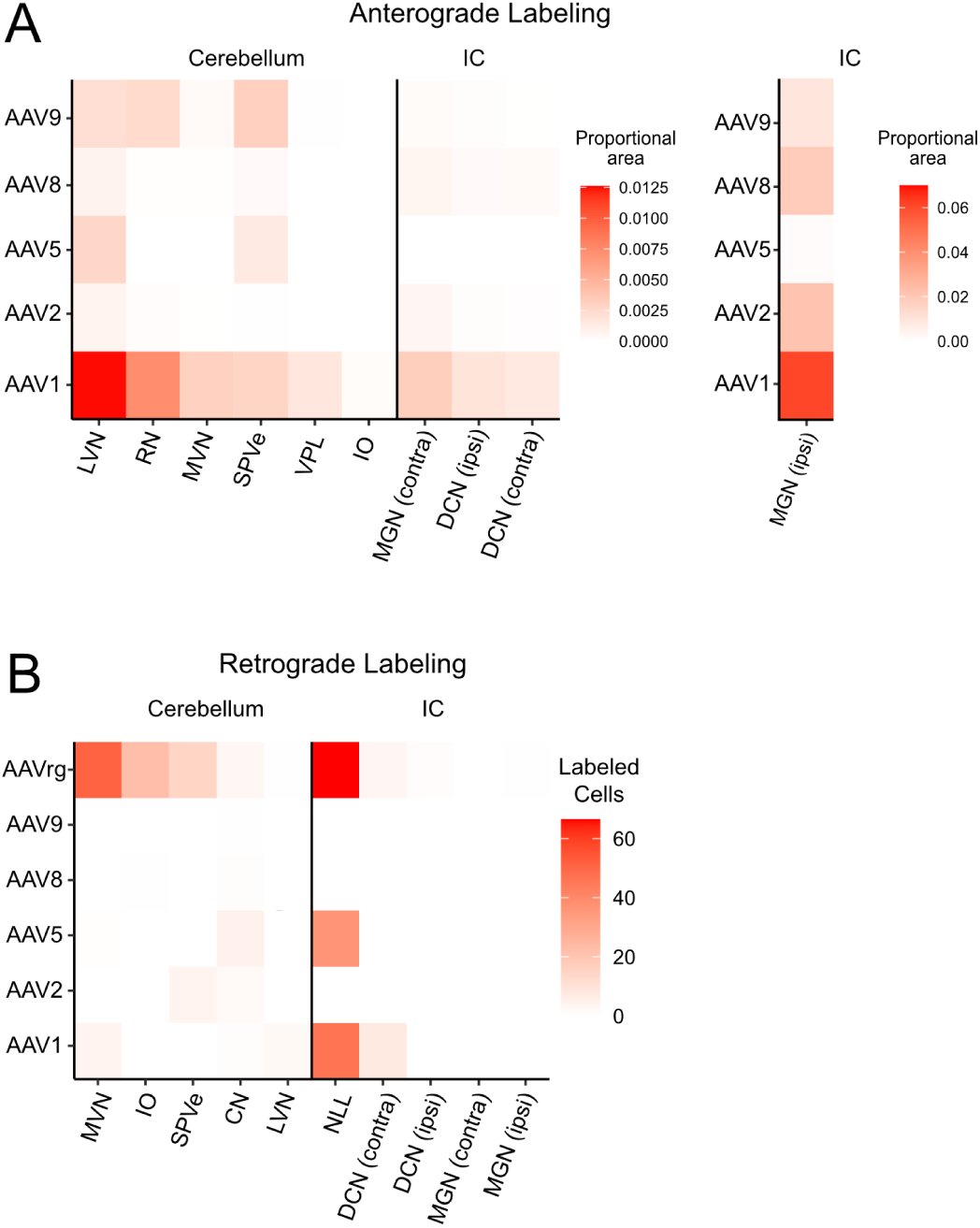
Summary of anterograde and retrograde labeling by serotype across regions of the midbrain and brainstem. Data from the previous figures were normalized to the total area of the nucleus and combined to create heatmaps of (A) anterograde and (B) retrograde labeling across serotypes and brain areas. Projections from IC to ipsilateral MGN labeled a much larger area than the other projections and are therefore plotted separately using a different color scale. AAV1 produced superior anterograde labeling compared to the other serotypes. As expected, AAVrg produced superior retrograde labeling. Overall, AAV8 and AAV9 had the lowest level of retrograde labeling, while AAV2 and 8 had the lowest anterograde labeling from the cerebellum and AAV5 had the lowest anterograde labeling from the IC.

## Discussion

AAVs are valuable tools to investigate neuronal function and connectivity and empirical studies are necessary to define their effectiveness in specific brain regions. Such studies allow researchers to choose the optimal serotype to address their experimental questions. Here we provide a systematic characterization of the efficacy of AAV serotypes in the cerebellum and IC, two regions important for multisensory integration and of interest to many researchers. First, we investigated the efficacy of AAV transduction in these regions by examining the extent of fluorescent labeling. Second, tropism within the cerebellum was evaluated by a careful analysis of labeled neurons, which was possible because of their distinct morphologies and localization. Next, differences in the extent of labeling of axonal projections in various targets of the IC and cerebellum were detailed. These differences are presumably due to variations in tropism or transduction efficacy of different populations of projection neurons. Finally, retrograde labeling was described across various input regions to the cerebellum and IC.

We found that AAV1 produces superior transgene expression defined by the number of neurons labeled, brightness of labeling, and volume of tissue labeled, in both the IC and in the cerebellum. Thus, AAV1 may be advantageous to transfect a large volume with a single injection in these regions. On the other hand, experiments requiring highly localized gene expression could benefit from the use of an alternate serotype. In the cerebellum, AAV1 transduced significantly more Purkinje cells and unipolar brush cells in lobule X of the cerebellar cortex compared to all other serotypes, and transduced significantly more molecular layer interneurons than AAV5, 9, and rg. Meanwhile, AAV2 transduced significantly more granule cells in lobule X of the cerebellar cortex than AAV1, 5, and 8. AAV2 was the most effective serotype for labeling granule cells. Although the labeling was rather dim in granule cells compared to other cell types, this level of gene expression is expected to be sufficient for Cre recombination and other processes that do not require strong overexpression of protein (Haery et al., 2019).

Labeled axonal projections were seen in a variety of IC and cerebellum output regions. Consistently, AAV1 produced the brightest axonal labeling compared to the other serotypes in the MGN, VPL, RN, MVN, ipsilateral DCN, ipsilateral MGN, IO, and LVN, which likely reflects higher transduction efficacy at the injection site or tropism for the specific projection neurons. AAV9 labeled a large number of projections in the red nucleus, which could indicate tropism of cerebellar nuclei neurons that project to this area (Low et al., 2018; Judd et al., 2021). The high degree of labeling in the MVN by AAV1 is likely due to superior transduction of Purkinje cells, whose axons project to MVN (Barmack, 2023).

Superior efficacy of AAV1 has been reported in other brain regions including the cortex, midbrain, corticospinal tract, nigrostriatal system, and red nucleus (Wang et al., 2003; Burger et al., 2004; McFarland et al., 2009; Blits et al., 2010; Hutson et al., 2012), while other studies demonstrate a lack of efficacy by AAV2 (Taymans et al., 2007; Aschauer et al., 2013; Watakabe et al., 2015) or higher efficacy of other serotypes (Paterna et al., 2004; Mason et al., 2010; Holehonnur et al., 2014). This region-specific variability of AAV efficacy underscores the necessity of studies such as this one.

All AAVs have been shown to cause some retrograde labeling in various brain regions (Taymans et al., 2007; Hollis Ii et al., 2008; Masamizu et al., 2011; Castle et al., 2014; Tervo et al., 2016; Sun et al., 2019; Haggerty et al., 2020). AAVrg was developed specifically to infect axons and retrogradely label neurons projecting to the area of injection (Tervo et al., 2016). In this study we found retrograde labeling of cells in NLL was higher for AAV1, 5, and rg than the other serotypes tested. AAV8 and 9 produced essentially no retrograde labeling in the regions analyzed, and may be useful serotypes when avoiding retrograde labeling is essential.

Enhancing the fluorescence that is present using an immunohistochemical approach would increase the number of neurons and processes that could be visualized, especially fine axonal processes that may have not have been detected here. This is only possible for anatomical studies that use fixed tissue. Higher expression may be necessary for physiological or behavioral studies that use AAVs to express optogenetic or chemogenetic tools or indicators of cellular activity. To increase expression, several modifications to the protocol used here could be made. The 2-week period between injection and analysis that we used is relatively short and longer periods may produce enhanced expression. The titer used here is lower than what is routinely available from commercial sources and a higher titer could increase expression. However, these modifications must be tested, as excessive expression caused by high titers or long incubation periods can cause toxicity (Watakabe et al., 2015).

In this study we only tested AAVs that used the hSyn promoter. A previous study showed that a CaMKII promotor can label Purkinje cells specifically and a GABAA receptor α6 subunit promotor can label granule cells specifically (Kim et al., 2015). Combining the GABAA receptor α6 subunit promotor with the AAV2 serotype may lead to higher expression in granule cells, which are often difficult to label effectively. The present study may provide guidance for neuroscientists planning studies that utilize AAVs to express transgenes in the IC and cerebellum.

## Acknowledgements

We thank Dr. Jason Newbern for confocal microscope use and Dr. Christopher Chen for helpful comments on the manuscript.

